# Structural insights into the RNA binding inhibitors of the C-terminal domain of the SARS-CoV-2 nucleocapsid

**DOI:** 10.1101/2024.10.11.617762

**Authors:** Preeti Dhaka, Jai Krishna Mahto, Ankur Singh, Pravindra Kumar, Shailly Tomar

## Abstract

The SARS-CoV-2 nucleocapsid (N) protein is an essential structural element of the virion, playing a crucial role in enclosing the viral genome into a ribonucleoprotein (RNP) assembly, as well as viral replication and transmission. The C-terminal domain of the N-protein (N-CTD) is essential for encapsidation, contributing to the stabilization of the RNP complex. In a previous study, three inhibitors (ceftriaxone, cefuroxime, and ampicillin) were screened for their potential to disrupt the RNA packaging process by targeting the N-protein. However, the binding efficacy, mechanism of RNA binding inhibition, and molecular insights of binding with N-CTD remain unclear. In this study, we evaluated the binding efficacy of these inhibitors using isothermal titration calorimetry (ITC), revealing the affinity of ceftriaxone (18 ± 1.3 μM), cefuroxime (55 ± 4.2 μM), and ampicillin (28 ± 1.2 μM) with the N-CTD. Further inhibition assay and fluorescence polarisation assay demonstrated RNA binding inhibition, with IC_50_ ranging from 10.4 to 12.4 μM and K_D_ values between 24 and 32 μM for the inhibitors. Additionally, we also determined the inhibitor-bound complex crystal structures of N-CTD-Ceftriaxone (2.0 Å) and N-CTD-Ampicillin (2.2 Å), along with the structure of apo N-CTD (1.4 Å). These crystal structures revealed previously unobserved interaction sites involving residues K261, K266, R293, Q294, and W301 at the oligomerization interface and the predicted RNA-binding region of N-CTD. These findings provide valuable molecular insights into the inhibition of N-CTD, highlighting its potential as an underexplored but promising target for the development of novel antiviral agents against coronaviruses.

**Highlights:**

- The inhibitors ceftriaxone, cefuroxime, and ampicillin-demonstrated high-affinity binding to the C-terminal domain (N-CTD) of the SARS-CoV-2 nucleocapsid (N) protein, effectively disrupting the formation of the N-CTD-RNA complex.
- Complex crystal structures of N-CTD with ceftriaxone and ampicillin revealed previously unobserved distinct binding sites.
- Structures reveal how the selected inhibitors disrupt the oligomerization of N-CTD and hinder the RNA packaging process of the virus.

## 1. Introduction

The severe acute respiratory syndrome coronavirus 2 (SARS-CoV-2) has caused the coronavirus disease 2019 (COVID-19) pandemic that led to many fatalities and economic consequences (Nicola et al., 2020). Despite widespread efforts, only a few licensed drugs are available to treat SARS-CoV-2 infections (Brady et al., 2024). Small molecules targeting viral proteins like RNA-dependent RNA polymerase (RdRp) protein hydroxychloroquine (Chakraborty et al., 2021; Eslami et al., 2020), lopinavir (Cao et al., 2020), remdesivir (Malin et al., 2020), and Nirmatrelvir (Owen et al., 2021) against proteases, have shown their potential inhibition against the SARS-CoV-2. Similarly, numerous drugs have been designed to target viral proteins directly in other viruses (Aggarwal et al., 2017; Fatma et al., 2020; Sharma et al., 2018). In addition to these compounds, Suramin (Boniardi et al., 2023) and Chicoric acid (Mercaldi et al., 2022) are proven promising antiviral compounds against the N-terminal domain (NTD) and C-terminal domain (CTD), respectively of SARS-CoV-2 nucleocapsid (N) protein. However, the available drugs have limitations like toxicity for different organs and potential incompatibility with other medicaments (Heskin et al., 2022) highlighting the necessity for the novel drug candidates with more effectiveness and fewer side effects.

The SARS-CoV-2 virus has a positive sense single-stranded RNA genome that codes for 16 non-structural proteins (nsps), including the proteases, Main proteases (Mpro), and papain-like proteases (PLpro) (Yan et al., 2022). Nsps are considered to play a role in viral replication, making them valuable targets for developing antivirals (Choudhary et al., 2022; Rani et al., 2022; Singh et al., 2023). In addition to the nsps, four structural proteins, including Spike (S), envelop (E), membrane (M) and the nucleocapsid (N) protein, also play crucial roles in viral genome protection and viral pathogenicity (Yan et al., 2022). Among structural proteins, the nucleocapsid (N) plays an essential role in packaging the viral RNA genome into a helical ribonucleoprotein (RNP) complex (Cubuk et al., 2021). The N-protein of SARS-CoV-2 protein is an approved antiviral drug targeting its evident involvement in the viral life cycle, including viral replication, transcription and encapsidation of the viral genome (McBride et al., 2014; Papageorgiou et al., 2016). N-protein helps in genome packaging and leads to the formation of ribonucleoprotein (RNP) complex on the establishment of a connection between viral RNA and the N-protein (McBride et al., 2014). The viral RNA interacts directly with the N-protein via its two functional domains, NTD and CTD, interconnected by highly disordered linker regions (Gordon et al., 2020; Morse et al., 2023). The earlier reported crystal structures of NTD and CTD have provided clear insight into the essential residues involved in the RNA encapsidation process, thereby regulating transcription (Sola et al., 2011) (Yang et al., 2021). The structural findings on CTD of N-protein (N-CTD) revealed its significant role in regulating viral replication and transcription after establishing specific contact with the transcriptional regulatory sequence (TRSs) of viral genome (Sola et al., 2015; Wu et al., 2014). The N-CTD’s critical role in recognizing conserved regulatory sequences in the SARS-CoV-2 genome and its involvement in the oligomerization process represents a promising target for developing inhibitors that can disrupt its oligomerization and RNA binding activities.

Recent structural studies have advanced our understanding of the N-CTD’s mechanism of RNA and small molecule binding (Mercaldi et al., 2022; Rafael Ciges-Tomas et al., 2022). For example, a crystal structure complex of N-CTD with chicoric acid has elucidated the inhibitory mechanism of this small molecule (Mercaldi et al., 2022). Considering the importance of this approach, it is crucial to explore new inhibitor binding sites on the N-CTD that can inhibit its oligomerization and RNA binding functions. Previously, three inhibitors, ceftriaxone, cefuroxime, and ampicillin, were screened as potential candidates for targeting the N-protein of SARS-CoV-2 using a high-throughput chip screening technique (Hu et al., 2021).

In this study, we first characterized the binding affinities of ceftriaxone, cefuroxime, and ampicillin to N-CTD using isothermal titration calorimetry (ITC). Subsequently, we employed a fluorescence intensity-based RNA binding assay and fluorescence polarisation (FP) assay to assess their impact on RNA binding activity. Furthermore, we determined the crystal structures of N-CTD in complex with ceftriaxone (2.0 Å) and ampicillin (2.2 Å) alongside the apo structure at high resolution (1.4 Å). The present findings reveal the molecular interactions between these inhibitors and the N-CTD protein, demonstrating their ability to hinder RNA binding and disrupt the packaging of the viral genome into the RNP complex.

## 2. Material and methods

### 2.1. Isothermal titration calorimetry (ITC)

ITC was used to determine the binding affinity of N-CTD protein with the selected inhibitors by analysing the dissociation constant (K_D_) and other thermodynamic parameters. These experiments were conducted using MicroCal ITC_200_ microcalorimeter (Malvern, Northampton, MA) at 25□°C and a stirring speed of 850 rpm. The purified N-CTD protein was dialyzed overnight at 4 °C in a phosphate buffer containing 20 mM sodium phosphate, 200 mM NaCl, and pH 8.0. Following dialysis, the protein and inhibitors were diluted to final concentrations of 20 μM and 500 μM, respectively. The buffer titrations against the N-CTD protein and buffer with inhibitors were performed as controls. Titrations consisted of 20 injections of 2µl of inhibitors into the cell solution containing N-CTD protein at a 10-fold lower concentration. After subtracting the mentioned controls, the isotherm curves were analysed using the analysis Software utilizing Origin7.0 (Edwards, 2002) associated with Microcal-ITC200 (commercially available). The output data was fitted to one-set site models to obtain non-linear curves and determined all the thermodynamic parameters, including the number of binding sites (n), association constant (K_A_), the binding enthalpy change (ΔH), binding entropy change (ΔS), and the equilibrium dissociation constant (K_D_) values.

### 2.2. RNA binding inhibition assay

A fluorescence intensity-based RNA binding assay was conducted to investigate the interaction of N-CTD protein with the RNA and estimate the inhibitory effect of the selected compounds (Byun et al., 2020; Kumar et al., 2021). To determine the potential of the inhibitors targeting the RNA-binding activity of N-CTD protein, the selected conserved genome (TRS) of SARS-CoV-2 was chemically synthesized with fluorescent 5’6-FAM-labelled 10-mer RNA probe (UCUCUAAACG) and obtained from Biotech Desk Pvt Ltd, India. The binding buffer includes 20 mM HEPES, pH 7.4, and 100 mM NaCl, which were then prepared using diethylpyrocarbonate (DEPC) treated RNAase-free water. The purified N-CTD protein was diluted in the binding buffer and added to each well in the range of 0-15 μM, followed by the addition of 1μM RNA in 96-well black flat-bottom Polystyrene microplates (Corning #3881). Incubation was carried out at 4°C for 30 minutes. Positive control wells with only 5’6-FAM-labelled RNA and negative control holding non-binding protein (Mpro) incubated with FAM-labelled RNA (Fig. 4a). Synergy HTX multimode plate reader (Agilent BioTek) included Gen5 software with a particular excitation wavelength (485/20nm)/emission wavelength of (528/20nm), and the read height of 8.5 mm, which was used to estimate the change in fluorescence intensity.

Further, the estimation of change in N-CTD-RNA interactions in the presence of an inhibitor was determined (Byun et al., 2020). The compounds were incubated overnight at 4°C with 15µM of purified N-CTD protein before observing the changes in interactions between the protein and FAM-RNA (1μM). Wells containing N-CTD protein incubated with FAM-RNA were used as positive controls, while wells with only FAM-RNA served as negative controls. After a 30-minute incubation, fluorescence intensities were measured. Graphs plotting percentage inhibition versus concentration were generated using GraphPad Prism 8.0, and the data represent the average of duplicate reactions.

### 2.3. Fluorescence polarisation (FP) assay

The chemically synthesized fluorescent 5’6-FAM-labelled RNA probe (UCUCUAAACG) of 10-mer was obtained from Biotech Desk Pvt Ltd (India). FP assays assisted in determining the binding affinity of the N-CTD protein with the RNA probe. The RNA probe was solubilized in a 20 mM HEPES buffer (pH 7.4) containing 100 mM NaCl, prepared from the stock solution for the experiment. All the buffers were prepared using DEPC-treated, autoclaved Milli-Q water to ensure an RNAase-free environment. The purified N-CTD protein with concentration gradient from 10 nM to 100 µM was diluted in a buffer composed of 50 mM sodium phosphate buffer (pH, 7.4) and 100 mM NaCl. Each protein concentration was thoroughly mixed with the RNA (20 µM as its final concentration) in 96-well flat bottom non-binding black (Greiner Bio-One) microplates. FP data acquisition was performed using Synergy H1 multimode plate reader (Agilent BioTek), with excitation (480 nm) and emission (535 nm) wavelengths. The acquired data was subjected to affinity binding curve analysis using the Hill1 model in OriginPro software.

After the affinity assessment of RNA for the CTD N-protein, FP analyses was conducted in the presence of selected inhibitor compounds (Lea and Simeonov, 2011; Mercaldi et al., 2022). Serial dilutions of the compound, ranging from 0.1 μM, to 500 μM, were prepared in a 50 mM sodium phosphate buffer (pH 7.4) with the addition of 100 mM NaCl. A concentration of 20 μM N-CTD protein was mixed with these serially diluted compounds and incubated for 60 min, followed by adding RNA at a concentration of 10 µM. FP assay measurements detected the binding affinity of the RNA probe to the N-CTD protein and the reduction of their interactions using inhibitor molecules. In the presence of inhibitors, N-CTD protein exhibited a reduced binding affinity for RNA, as reflected by the corresponding K_D_ values. The error bars represent the standard deviation (SD) from the means of duplicate reactions.

### 2.4. Crystallization and data collection

N-CTD of SARS-CoV-2 N-protein was concentrated up to 22 mg/ml and subjected to crystallization after incubation with the selected inhibitors for 1 hour using the sitting-drop vapour diffusion method at 20 °C. Best diffracting crystals were obtained from the crystal buffer with 0.2 M Lithium sulphate, 0.1 M Tris (pH 7.8) and 31% polyethylene glycol (PEG) 4000 compositions. Crystals of N-CTD protein and inhibitors (ceftriaxone and ampicillin) complexes were obtained after mixing 1mM of the inhibitor into the 22mg/ml protein. Crystals were cryo-protected in a reservoir solution containing 10% ethylene glycol, was used as cryo-protectant for the N-CTD protein crystals during the data collection, and then flash-frozen using nitrogen stream (100 K). Diffraction data for N-CTD protein complexes were collected at the Home Source, Macromolecular Crystallography Unit (MCU) facility, and Institute Instrumentation Centre (IIC) at the Indian Institute of Technology Roorkee (IIT) Roorkee.

### 2.5. Structure determination and refinement

The crystal data from X-ray diffraction were collected at the beamline using a wavelength of 1.4 Å (apo N-CTD) 2.0 Å (CTD-Ceftriaxone) and 2.2 Å (CTD-ampicillin). The structure determination was performed with the molecular replacement method (MR) (Panjikar et al., 2009), employing the X-ray crystal structure of SARS-CoV-2 N-CTD protein (PDB ID: 6YUN) as a search model (Zinzula et al., 2021). All initial data were further refined with Refmac5 (Murshudov et al., 2011) as part of the CCP4i2 program (“The CCP4 suite: Programs for protein crystallography,” 1994) and interactively adjusted by COOT (Emsley et al., 2010). The final statistics for the X-ray diffraction and the model refinement are listed in Table 2.

## 3. Results

### 3.1. ITC of inhibitors binding to N-CTD

The binding affinities of ceftriaxone, cefuroxime, and ampicillin with the N-CTD were quantified using isothermal titration calorimetry (ITC), which revealed the thermodynamic signatures of binding, including K_D_ values as well as enthalpic and entropic contributions (Table 1). The K_D_ values for ceftriaxone (18 ± 1.3 μM), cefuroxime (55 ± 4.2 μM), and ampicillin (28 ± 1.2 μM) are significantly lower than that of N-CTD’s interaction with RNA (16 ± 1.4 μM) (Fig. 1). Notably, the consistent entropic contributions in the binding of these inhibitors suggest similar conformational changes in the N-CTD upon binding. Overall, these results highlight the high-affinity interactions between the inhibitors and N-CTD.

**Fig. 1.**
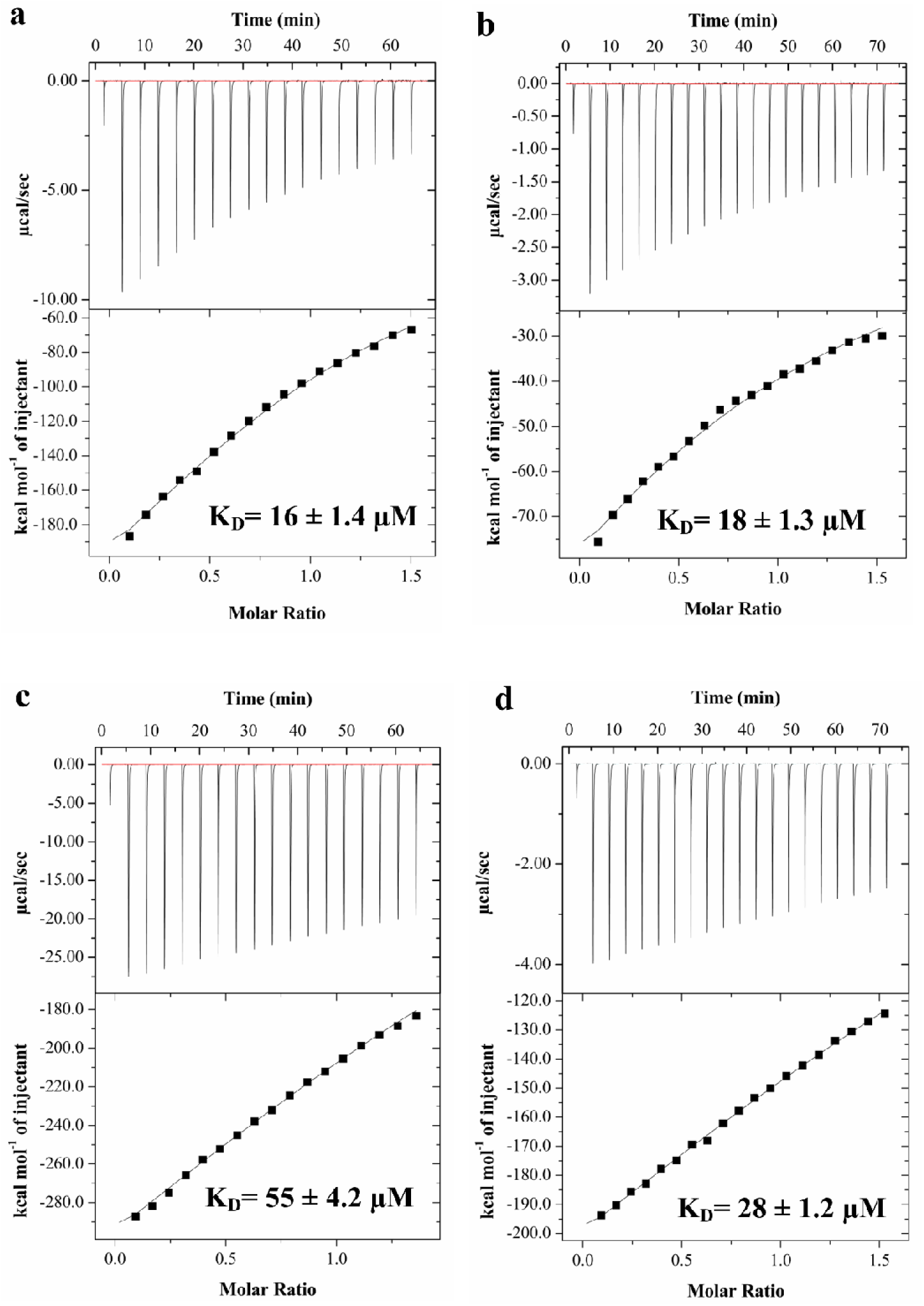
Thermodynamic analysis of N-CTD interactions with RNA and the inhibitors using ITC. Isothermal titration calorimetry (ITC) profiles showing the binding interactions between N-CTD (20 μM) and (a) RNA (200 μM), (b) ceftriaxone (500 μM), (c) cefuroxime (500 μM), and (d) ampicillin (500 μM).

**Table 1.**
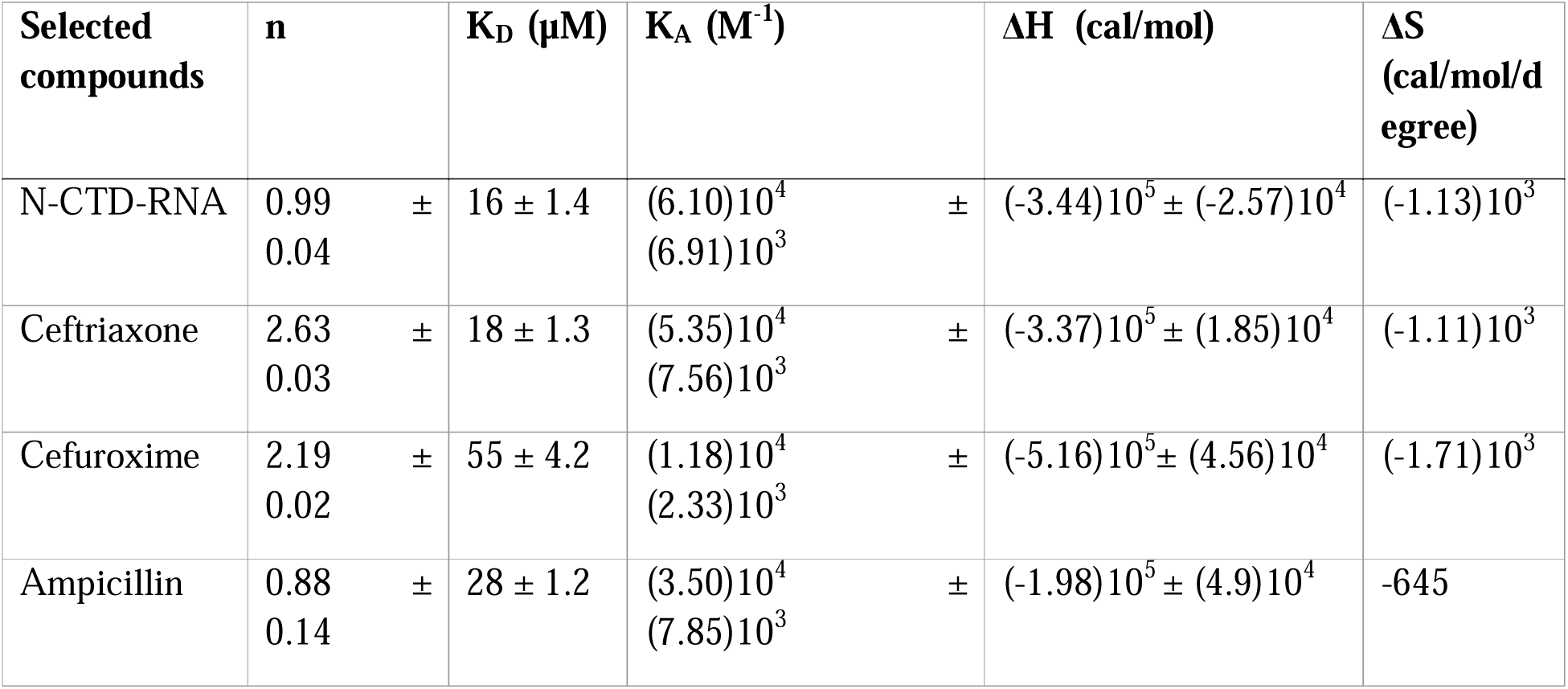
The thermodynamic isotherm parameters were obtained from biophysical interaction analysis of compounds with N-CTD protein using the ITC.

### 3.2. RNA binding inhibition assay

A fluorescence intensity-based assay was employed to assess the inhibitory potential of the inhibitors on the interactions between the N-CTD and FAM-labelled RNA. The fluorescence intensity increased proportionally with the concentration of N-CTD, reaching saturation at 15 µM concentration of N-CTD (Fig. 2a). In the presence of inhibitors, a significant reduction in fluorescence intensity was observed when incubated with a saturating concentration of N-CTD, indicating inhibition of N-CTD-RNA complex formation. Using GraphPad Prism (version 8.0), the half-maximal inhibitory concentration (IC_50_) values were determined for ceftriaxone (IC_50_=12.4 ± 2.4 µM), cefuroxime (IC_50_ =15.4 ± 2.8 µM), and ampicillin (IC_50_=10.4 ± 0.2 µM) (Fig. 2b-d). These selected compounds are shown to disturb the N-CTD interactions with the RNA efficiently.

**Fig. 2.**
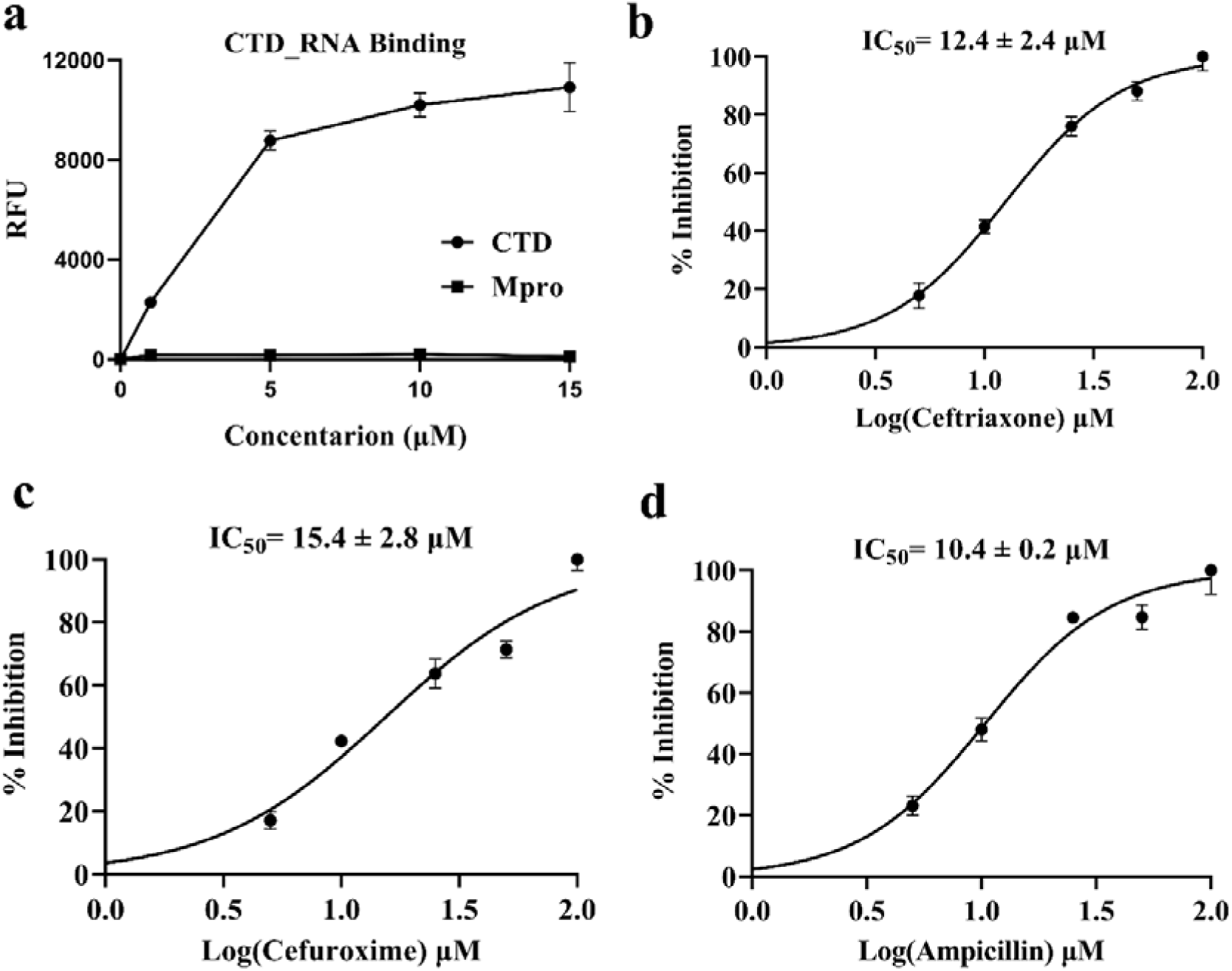
Fluorescence-based RNA binding and inhibition assay. (a) Fluorescence intensity changes of FAM-labeled RNA upon binding to the N-CTD protein, demonstrating the N-CTD-RNA interaction. The dose-response curves show the percentage inhibition of N-CTD-RNA binding by (b) ceftriaxone, (c) Cefuroxime, and (d) ampicillin, plotted against varying inhibitor concentrations. Graphs were generated using GraphPad Prism (version 8.0) with error bars representing the standard deviation from duplicate measurements

### 3.3. Fluorescence polarisation (FP) assay

Fluorescence polarisation (FP) concentration-response sequel experiments were conducted with the inhibitors further to validate the inhibition of N-CTD and RNA interactions. Initially, N-CTD was titrated against RNA to establish the saturation of the fluorescence polarisation signal, which was achieved at N-CTD concentration of 20 µM (Fig. 3a). At this saturating N-CTD concentration, ceftriaxone, cefuroxime, and ampicillin were titrated, resulting in a noticeable decline in the fluorescence signal, indicating disruption of N-CTD-RNA complex (Fig. 3b-d). The K_D_ values for ceftriaxone (32 ± 6.4 µM), cefuroxime (29 ± 3.9 µM), and ampicillin (24 ± 4.1 µM) demonstrate that these inhibitors effectively disrupt the interactions of N-CTD and RNA.

**Fig. 3.**
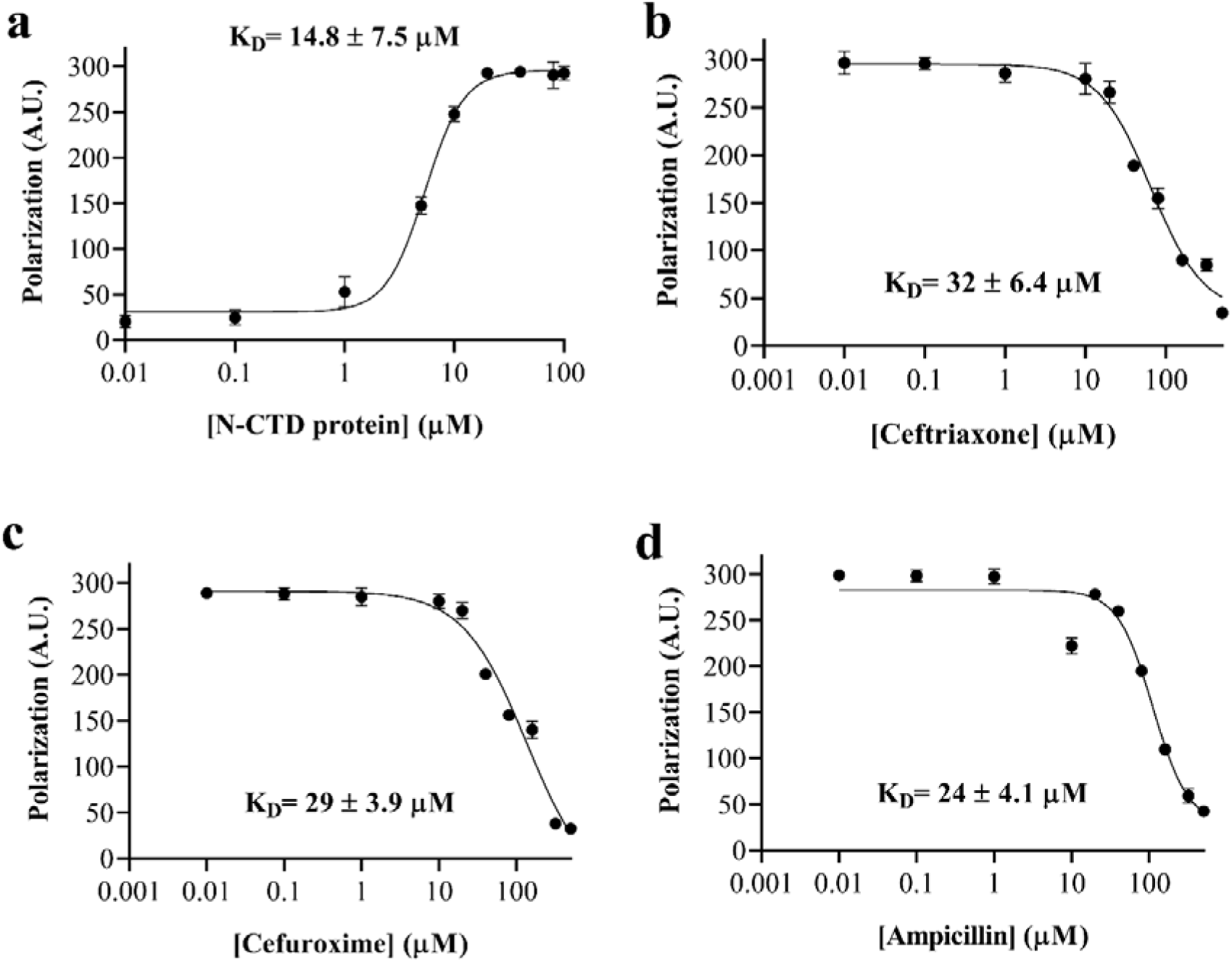
Fluorescence polarisation assay for N-CTD-RNA interaction and inhibition. (a) FP signal from 5’6-FAM-labelled RNA shows a concentration-dependent increase in polarization upon binding to N-CTD. FP signal response illustrating the inhibitory effects of increasing concentrations of (b) ceftriaxone, (c) cefuroxime, and (d) ampicillin on N-CTD-RNA binding. Error bars represent the standard deviation (SD) derived from the means of two independent experiments.

### 3.4. N-CTD crystal structures

The crystal structure of the N-CTD protein in complex with ceftriaxone and ampicillin compounds were determined to elucidate their binding modes and mechanisms of RNA-binding inhibition. The crystal structures of the N-CTD apo form, as well as N-CTD bound to ceftriaxone and ampicillin, were refined to resolutions of 1.4, 2.0 and 2.2 Å, respectively (Table 2). The phases were obtained using the molecular replacement method with previously solved N-CTD structure (PDB ID: 6YUN) as the search model. Detailed data collection statistics are given in Table 2.

**Table 2.** Data collection and refinement statistics.

| Property | N-CTD protein (Apo) | N-CTD protein in complex with ceftriaxone | N-CTD protein in complex with ampicillin |
| --- | --- | --- | --- |
| **Data collection statistics** |  |  |  |
| **Resolution range (Å)** | 24.0-1.4 (1.5-1.4) | 26.6-2.0 (2.07-2.0) | 24.1-2.2 (2.9-2.0) |
| **Space group** | *P1 2<sub>1</sub> 1* | *P1* | *P1 2<sub>1</sub> 1* |
| **Unit cell** | 71.0 43.5 74.8<br>90.0 94.5 90.0 | 43.5 46.6 69.0<br>106.6 90.1 93.5 | 44.6 43.6 56.7<br>90.0 108.7 90.0 |
| **Unique reflections** | 84966 (7977) | 34800 (3450) | 10631 (1061) |
| **Multiplicity** | 3.7 (2.4) | 4.2 (2.7) | 5.5 (5.8) |
| **Completeness (%)** | 94.4 (89.2) | 99.2 (98.6) | 99.6 (99.9) |
| **Mean I/sigma(I)** | 17.2 (2.8) | 3.22 (1.9) | 9.17 (2.2) |
| **Wilson B-factor** | 8.0 | 15.1 | 12.5 |
| **R-merge** | 0.04 (0.31) | 0.06 (0.27) | 0.09 (0.15) |
| **R-means** | 0.05 (0.39) | 0.07 (0.32) | 0.09 (0.16) |
| **R-pim** | 0.02 (0.23) | 0.03 (0.18) | 0.04 (0.07) |
| **CC1/2** | 0.99 (0.83) | 0.99 (0.88) | 0.99 (0.98) |
| **Refinement statistics** |  |  |  |
| **R-factor<sup>a</sup>** | 0.14 | 0.16 | 0.19 |
| **R-free<sup>b</sup>** | 0.17 | 0.21 | 0.25 |
| **No. of atoms** | 4504 | 4417 | 1993 |
| **macromolecules** | 3673 | 3628 | 1735 |
| **ligands** | 0 | 120 | 28 |
| **solvent** | 831 | 669 | 230 |
| **Protein residues** | 445 | 445 | 216 |
| **r.m.s.d bond lengths (Å)** | 0.01 | 0.01 | 0.01 |
| **r.m.s.d bond angles (°)** | 1.9 | 1.6 | 1.7 |
| **Ramachandran favored (%)** | 98.8 | 99.0 | 98.1 |
| **Ramachandran allowed (%)** | 1.1 | 0.9 | 1.9 |
| **Ramachandran outliers (%)** | 0.0 | 0.0 | 0.0 |
| **Rotamer outliers (%)** | 2.3 | 1.6 | 3.9 |
| **Clashscore<sup>c</sup>** | 2.6 | 5.6 | 8.3 |
| **Average B-factor (Å<sup>2</sup>)** | 13.5 | 20.6 | 15.3 |
| **macromolecules** | 10.1 | 17.6 | 13.8 |
| **ligands** | 0 | 43.8 | 38.4 |
| **solvent** | 28.2 | 32.4 | 23.1 |

<sup>a</sup>Rfactor = $\sum hkl |F_o(hkl) - F_c(hkl)| / \sum hkl F_o(hkl)$ . <sup>b</sup>R<sub>free</sub> was calculated for a test set of reflections (5.0 %) omitted from the refinement.
<sup>c</sup>Clashscore is defined as the number of clashes calculated for the model per 1000 atoms (including hydrogens) of the model.

Consistent with previously reported structures (Jia et al., 2022; Luan et al., 2022; Peng et al., 2020; Rafael Ciges-Tomas et al., 2022; Yang et al., 2021; Ye et al., 2021, 2020; Zinzula et al., 2021) the N-CTD protomer feature five α-helices and three η helices, along with the two β antiparallel-strands (Fig. 4a, 4d). The N-CTD dimer is primarily stabilized by hydrogen bonding interactions between β2 strands (G328–I337) of each protomer (Zhou et al., 2020). Additional stabilization comes from interactions involving residues in the loop connecting α1 and α2 helices (R277, G278, E280, Q283 and N285) and residues within the α4 helix and β1 strand (G316, S318, R319, and I320).

**Fig. 4.**
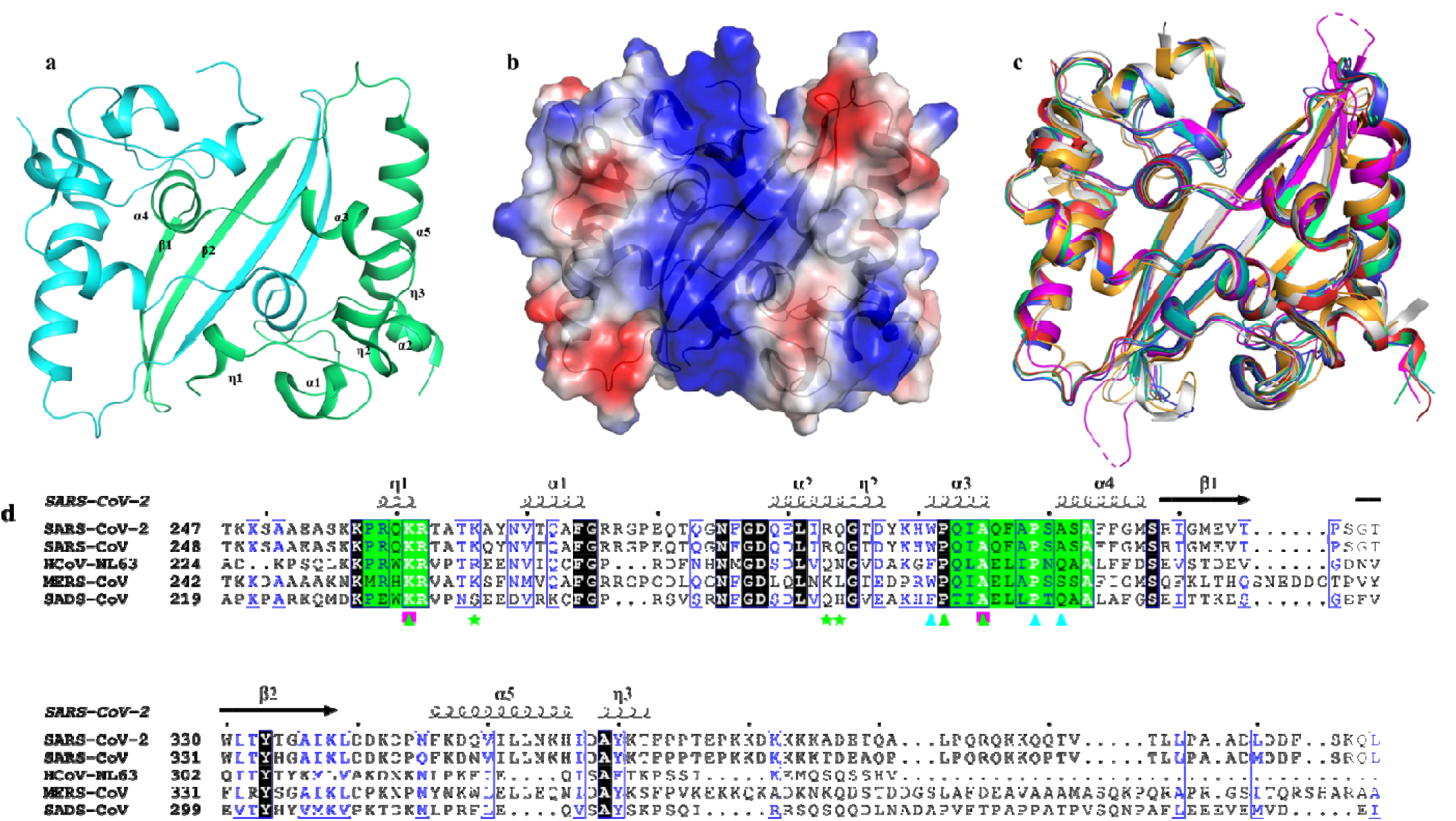
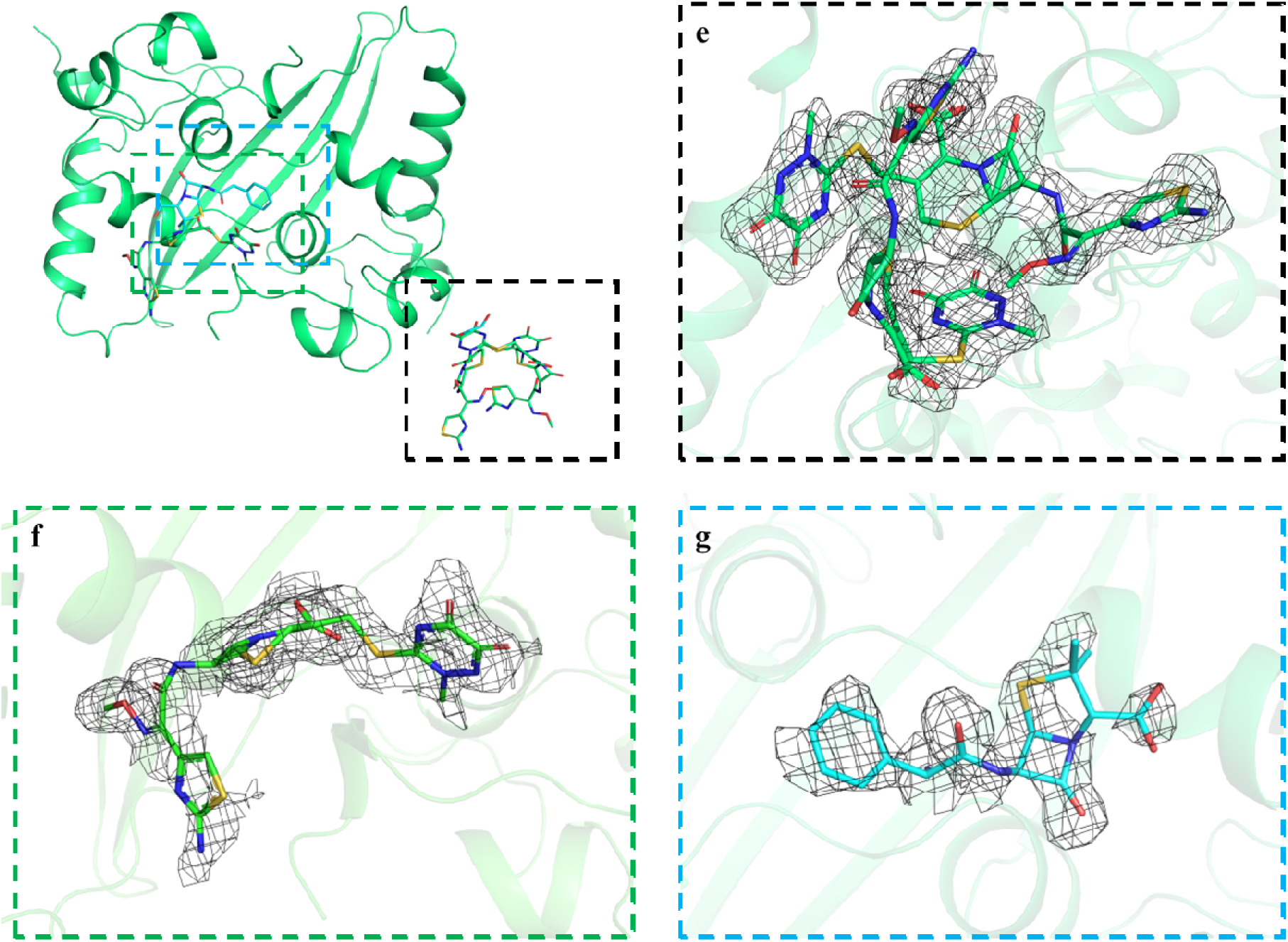
Crystal structure of N-CTD and inhibitor complexes. (a) Cartoon representation of the N-CTD dimer, with two protomers colored in green and cyan. Each protomer has five α-helices, three η-helices, and two β-strands. (b) A positively charged groove is formed at the helix face by η1, α3-helices, α4-helices. (c) Structural superposition of the N-CTD dimer from SARS-CoV-2 (green) with related structures: SARS-CoV-2 (PDB ID: 7CE0; teal), SARS-CoV (PDB ID: 2GIB; red), SARS-CoV (PDB ID: 2CJR; blue), MERS-CoV (PDB ID: 6G13; magenta), SADS-CoV (PDB ID: 8YM1; orange), HCoV-NL63 (PDB ID: 5EPW; blue). (d) Sequence alignment of N proteins from six related coronaviruses. The secondary structure elements α□helix, η□helix, and β□strand are shown. Identical residues are highlighted in black background, and similar residues are labelled in blue and boxed in blue lines. Green star, green triangle, cyan triangle, and purple square denotes residues interacting to ceftriaxone site first, ceftriaxone site second, ampicillin, and cefuroxime, respectively. The green background highlights residues that form the positively charged groove of N-CTD. The program ESPript was used for visualization. Electron density omit maps (contoured at 2.5σ) for ceftriaxone binding at site (d) first and (e) second. (f) Omit map (contoured at 2.5σ) showing ampicillin binding. Figures were prepared using PyMol (Robert and Gouet, 2014)

In the apo form, a positively charged groove is present on the helix face, formed by residues from the η1, α3-helices, α4-helices and loops connecting α3 and α4 helices, which is predicted to function as the RNA binding groove (Fig. 4b). Superposition of N□CTD structure with the N□CTD structures of SARS-CoV-2 virus (Zhang et al., 2024), SARS□CoV (Chen et al., 2007), MERS□CoV (Nguyen et al., 2019), HCoV□NL63 (Szelazek et al., 2017), and SADS-CoV (Zhang et al., 2024), yielded an RMSD of 0.30–1.46 Å (Fig. 4c). Interestingly, the positively charged groove forming residues are conserved in all these structures (Fig. 4d), supporting its role as the RNA binding site in these viruses.

In the N-CTD complexes with ceftriaxone and ampicillin, both inhibitors show a good fit to the electron density (Fig. 4e-g). However, the β-lactam ring of ampicillin showed poorer density, possibly due to the degradation of the b-lactam ring under crystallization conditions (Siddiqui et al., 2014). Three ceftriaxone molecules were observed binding to the N-CTD dimer. Two of these molecules form a ball-like cluster, and interact with the surface of one N-CTD protomer, while the third binds at the positively charged groove present on the helix face. The ampicillin molecule was also observed to bind at this pocket near the ceftriaxone binding site.

In contrast to the binding of chicoric acid molecule (Mercaldi et al., 2022), the N-CTD complexes with ceftriaxone and ampicillin exhibit distinct interaction sites. The first ceftriaxone binding site, adjacent to the α3 helix, is characterized by ionic interactions with R293 and Q294 residues, along with hydrogen bonding interaction between K266 and the carbonyl oxygens and nitrogen atoms of the cyclic 1,2-diamide moiety of the ceftriaxone molecule (Fig. 5b). A second ceftriaxone molecule is stabilized through van der Waals forces, positioning its 7-aminocephalosporanic acid core sandwiched between cyclic 1,2-diamide and β-lactam carbonyl of the first molecule (Fig. 5b). Furthermore, the 7-aminocephalosporanic acid cores of both bound ceftriaxone molecules interact via van der Waals forces, creating a ball-like cluster. The second binding site, proximal to the α4 helix, involves ionic interactions with residue K261 and hydrogen bonding interaction with the main chain of P302 and A305 residues (Fig. 5c).

**Fig. 5.**
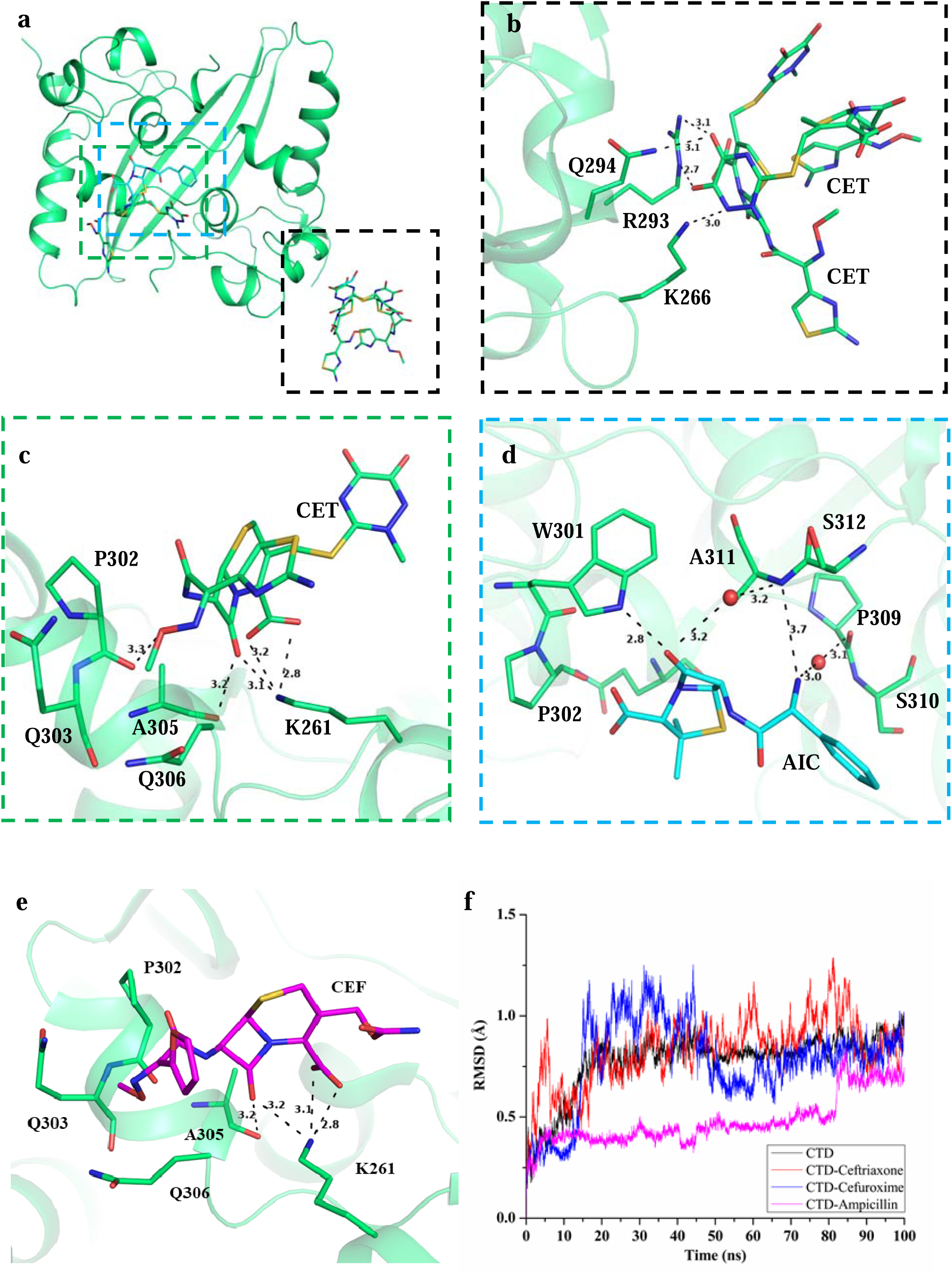
Structural analysis of N-CTD complexes with ceftriaxone, ampicillin, and cefuroxime. (a) Cartoon representation of the N-CTD dimer bound with ceftriaxone and ampicillin. (b) The ceftriaxone cluster is anchored to the N-CTD surface via interactions that are highlighted here. (c) A ceftriaxone molecule (CET) is positioned within the positively charged groove via interactions that are highlighted here. (d) Similarly, an ampicillin (AIC) molecule is anchored within the positively charged groove via interactions that are shown here. (e) The docked conformation of cefuroxime (CEF) molecule. (f) The RMSD plots from the molecular dynamics (MD) simulations of N-CTD complexes with ceftriaxone, cefuroxime, and ampicillin, providing insights into their binding stability. The interactions are represented with black dotted lines. Residues and bond distances (in Å) are labelled.

In the N-CTD-ampicillin complex, ampicillin binds proximal to the α4 helix as well, close to the ceftriaxone binding site. The binding of ampicillin to the N-CTD is stabilized by ionic interactions involving residue W301 and hydrogen bonding interaction with the main chain of A311 (Fig. 5d). Additionally, ampicillin interacts with two water molecules, coordinated by main chain of A311 and P309 residues.

Attempts to obtain a crystal structure of the N-CTD complex with cefuroxime were unsuccessful, likely due to the very short half-life of cefuroxime molecule. As an alternative, the cefuroxime molecule was docked at the ceftriaxone binding site because of its structural analogy to ceftriaxone molecule. The cefuroxime bound N-CTD structure exhibited a similar RMSD profile to the ceftriaxone bound N-CTD structure when simulated for 100 ns (Fig. 5f). The binding pose of cefuroxime is characterized by ionic interactions with K261, and hydrogen-bonding interaction with the main chain of A305 residue, closely resembling the interactions observed with ceftriaxone (Fig. 5e).

A previous study (Chen et al., 2007) proposed a model for N-CTD oligomerization and RNA encapsidation, featuring a helical arrangement of N-CTD dimers that form an X-shaped octamer (Fig. 6c). This structural organization creates a positively charged groove that facilitates the helical wrapping of RNA, as illustrated in Fig. 6d. Within this X-shaped arrangement, a critical interaction between dimers involve α2 and η1 helical residues, where ionic interactions with residue K261, R293 and Q294 play a pivotal role (Fig. 6e-f). Notably, the ceftriaxone cluster binds at this same interface, suggesting that ceftriaxone disrupts N-CTD oligomerization by interfering with these critical interactions (Fig. 6g). The conservation of these dimer interface residues between SARS-CoV-2 and SARS-CoV (Fig. 4d) further supports the potential for ceftriaxone to disrupt oligomerization in both viruses.

**Fig. 6.**
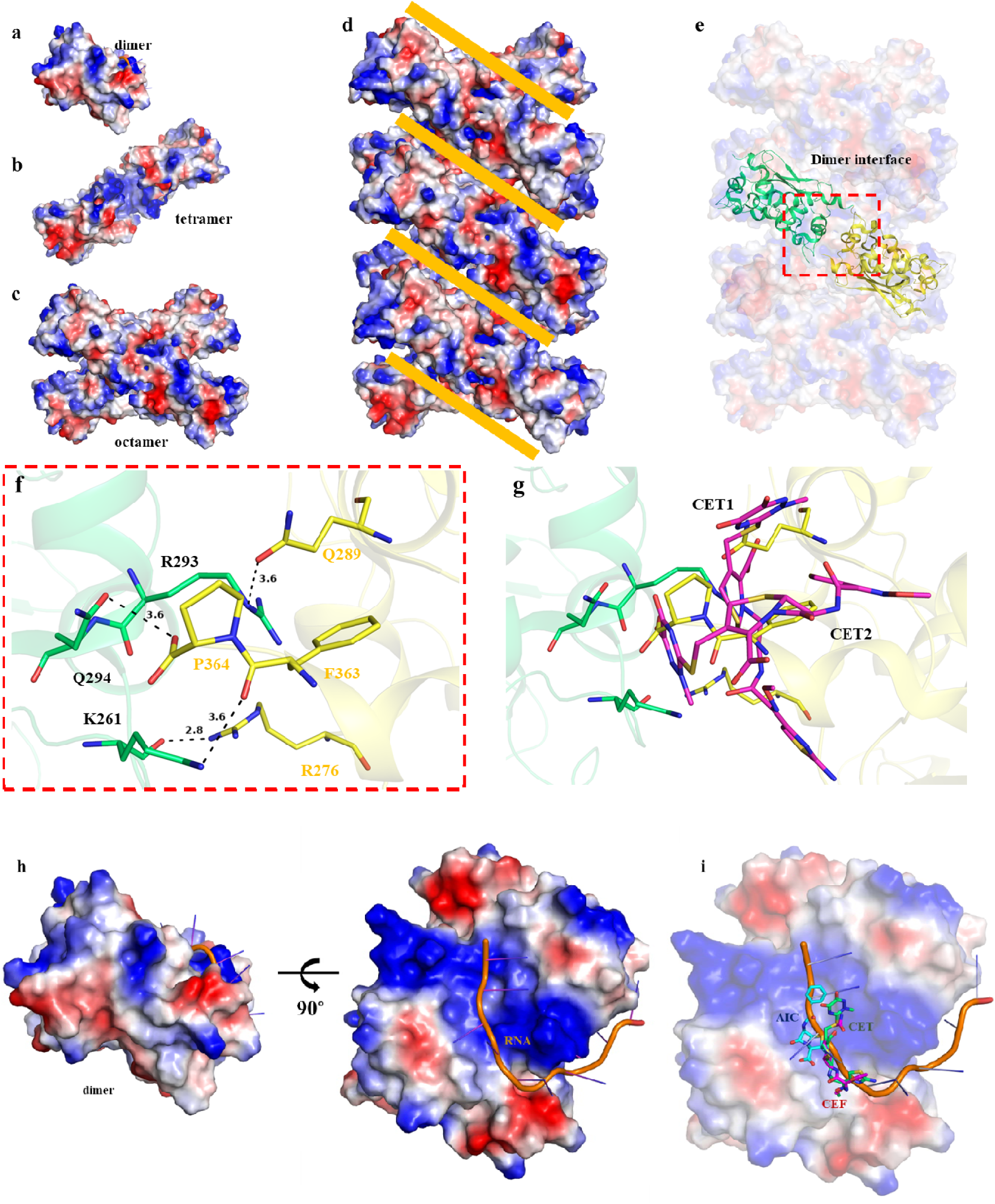
Inhibition of N-CTD oligomerization and RNA encapsidation by ceftriaxone, cefuroxime, and ampicillin. (a) N-CTD dimers are arranged into (b) a tetramer and subsequently to (c) a X-shaped octamer structure (d) RNA helically wraps around the X-shaped octamers. (e) The X-shaped octamer is stabilized by interactions between two dimers forming a tetramer. (f) Key interactions between the dimers that drive oligomerization are highlighted. (g) Ceftriaxone (CET1 and CET2) binding at site first on N-CTD surface disrupts these crucial interactions, thereby inhibiting the oligomerization process. (h) An Alphafold3 predicted model of RNA binding to N-CTD shows binding within the positively charged groove formed at the helix face of the N-CTD dimer. (i) Ceftriaxone (CET), cefuroxime (CEF), and ampicillin (AIC) interfere with RNA binding to this groove, thus disrupting the RNA encapsidation process.

Finally, to assess the binding consequences of these three inhibitor molecules to the RNA binding, an N-CTD RNA complex was generated with Alphafold3 (Abramson et al., 2024). The predicted model shows that the RNA binds in the positively charged groove at the helix face of the dimer (Fig. 6h). Interestingly, the same groove has been observed to bind ceftriaxone, ampicillin as well as cefuroxime, suggesting these inhibitors directly inhibit the RNA binding to N-CTD (Fig. 6i). Interacting residues to these inhibitors at the positively charged groove are conserved across *coronaviridae* family suggesting (Fig. 4d), these molecules could inhibit RNA encapsidation in all related viruses.

## 4. Discussion

Identifying and characterizing small molecule binding sites in target proteins through a crystallographic approach has been highly beneficial for drug discovery and scaffold optimization (Caines et al., 2012; Günther et al., 2021; Narayanan et al., 2022; Rochel et al., 2000; Sabini et al., 2003). In line with this approach, three inhibitors—ceftriaxone, cefuroxime, and ampicillin—recently screened against the N-protein of SARS-CoV-2 (Hu et al., 2021) were investigated through crystallographic studies for their binding to the N-CTD. These compounds have been previously used to treat intra-abdominal and respiratory tract infections, and ceftriaxone is currently under Recruiting Phase 3 Trials for community-acquired pneumonia, influenza, and COVID-19 treatment (Angus et al., 2020; Blum et al., 2023; Sturza et al., 2023). Belonging to the cephalosporin class, these drugs are considered promising repurposing candidates targeting the N-CTD (Hu et al., 2021; Luan et al., 2022). Their proposed mechanism disrupts N-protein oligomerization, hindering the virus’s RNA packaging process. However, there remains a knowledge gap in understanding their efficiency in binding to the N-CTD and their ability to interfere with the RNA binding. Moreover, the exact binding mode of these inhibitors to the N-CTD protein was unknown. To address these gaps, biochemical and biophysical assays were performed to assess their binding affinity and inhibitory potential on RNA binding. Crystallographic studies further elucidated the binding modes of ceftriaxone and ampicillin inhibitors with the N-CTD.

The binding affinities of the RNA, ceftriaxone, cefuroxime, and ampicillin against the N-CTD protein were assessed using ITC, revealing that N-CTD exhibits a good affinity for the inhibitors, with K_D_ ranging from 18 to 55 µM. Furthermore, both the fluorescence-based RNA binding assay and the fluorescence polarization assay indicated that the inhibitors impede RNA binding with K_D_ values between 24-32 µM for these inhibitors. These results are consistent with a previous study that reported a similar K_D_ of 41 µM for RNA binding inhibition by chicoric acid (Mercaldi et al., 2022). However, chicoric acid itself demonstrates a significantly higher binding affinity to N-CTD, with K_D_ value of 0.25 µM, suggesting a multiple-fold higher binding affinity with N-CTD, though the RNA inhibition profiles are comparable.

Contrary to the predicted site, the crystal structures of the N-CTD inhibitor complexes reveal distinct binding locations for the inhibitors. A previous *in silico* study predicted that residues D341, L339, and R259 were responsible for the ceftriaxone binding to N-CTD (Hu et al., 2021). However, our crystallographic and structural modelling data show that ceftriaxone, ampicillin and cefuroxime bind to different sites, with K261, K266, R293, Q294, and W301 playing key roles in binding rather than the predicted residues. Interestingly, these interacting residues are seen to bind RNA in the Alphafold3 model of N-CTD and RNA complex, suggesting that these inhibitors disrupt the RNA encapsidation process by binding to these critical residues.

The crystal structure of N-CTD in complex with ceftriaxone further reveals a dual inhibitory mechanism, as it binds at two distinct sites. First, at the surface, ceftriaxone disrupts N-CTD oligomerization, preventing the proper assembly of the nucleocapsid. Second, at the RNA binding groove, ceftriaxone impedes the RNA binding. Previous studies have majorly emphasized on targeting the dimerization site and RNA binding site of N-Protein (Cubuk et al., 2021; Dang and Song, 2022; Luan et al., 2022; Mercaldi et al., 2022; Peng et al., 2020; Rafael Ciges-Tomas et al., 2022; Wei et al., 2024). Additionally, crystallographic study on chicroic acid N-CTD complex has reinforced this approach (Mercaldi et al., 2022). In line with this approach, the molecule 5-Benzyloxygramine, previously identified as an inhibitor of N protein oligomerization (Lin et al., 2020), was investigated with a crystal structure complex with N-NTD (Bhutkar et al., 2022; Hong et al., 2024). This study elucidated its N-NTD binding mechanism and its ability to induce non-native dimerization of N-NTD (Hong et al., 2024). While studies have largely focused on N-NTD (Dhaka et al., 2023; Kumari et al., 2024; Luan et al., 2022; Peng et al., 2020; Wang et al., 2022; Wei et al., 2024), N-CTD dimerization has been underexplored as a target for the discovery of antivirals. Our results demonstrate that ceftriaxone inhibits N-CTD dimerization, providing a new avenue for therapeutic intervention against SARS-CoV-2 and related viruses.

In conclusion, our findings demonstrate that ceftriaxone, cefuroxime and ampicillin exhibit strong binding affinity to N-CTD and effectively disrupt RNA binding. The crystallographic structures provide mechanistic insights, revealing previously unrecognized inhibitor binding sites binding sites that interfere with both N-CTD oligomerization and RNA encapsidation. This study underscores the potential of these small molecules (ceftriaxone, cefuroxime, and ampicillin) as inhibitors of N-CTD, offering a promising avenue for targeting essential structural components of SARS-CoV-2 and related viruses to hinder viral replication.

## CRediT authorship Contribution statement

Conceptualization, P.K., and S.T.; Methodology, P.K., S.T., P.D., J.K.M., A.S., and K.J., S.V.; Experimentation, P.D., J.K.M., and A.S.; Formal analysis, P.K., and S.T.; Writing-original draft, P.D., J.K.M., and A.S.; Writing-review & editing, P.K., and S.T., Supervision, P.K., and S.T.; All authors read, revised and approved the manuscript.

## Data availability

The crystal structures generated during the current study are available in the Protein Data Bank (PDB) under the accession numbers, 9IN1 (N-CTD Apo), 8ZFV (N-CTD-ceftriaxone) and 8W6W (N-CTD-ampicillin), and through the links: https://www.rcsb.org/structure/9IN1, https://www.rcsb.org/structure/8ZFV, and https://www.rcsb.org/structure/8W6W, respectively. The authors have included the data in the materials and methods and the result section of the manuscript.

## Acknowledgements

The authors thank the Department of Biosciences and Bioengineering (BSBE) for providing the lab facility, Bioinformatics Centre (BIC), supported by the Government of India (reference number BT/PR40141/BTIS/137/16/2021), the Macromolecular Crystallographic Facility (MCU) for the computer facility and Ashok Soota Molecular Medicine Facility, BSBE, Indian Institute of Technology Roorkee (IIT Roorkee).

## References

Abramson, J., Adler, J., Dunger, J., Evans, R., Green, T., Pritzel, A., Ronneberger, O., Willmore, L., Ballard, A.J., Bambrick, J., Bodenstein, S.W., Evans, D.A., Hung, C.-C., O’Neill, M., Reiman, D., Tunyasuvunakool, K., Wu, Z., Žemgulytė, A., Arvaniti, E., Beattie, C., Bertolli, O., Bridgland, A., Cherepanov, A., Congreve, M., Cowen-Rivers, A.I., Cowie, A., Figurnov, M., Fuchs, F.B., Gladman, H., Jain, R., Khan, Y.A., Low, C.M.R., Perlin, K., Potapenko, A., Savy, P., Singh, S., Stecula, A., Thillaisundaram, A., Tong, C., Yakneen, S., Zhong, E.D., Zielinski, M., Žídek, A., Bapst, V., Kohli, P., Jaderberg, M., Hassabis, D., Jumper, J.M., 2024. Accurate structure prediction of biomolecular interactions with AlphaFold 3. Nature 630, 493–500. 10.1038/s41586-024-07487-w

Aggarwal, M., Kaur, R., Saha, A., Mudgal, R., Yadav, R., Dash, P.K., Parida, M., Kumar, P., Tomar, S., 2017. Evaluation of antiviral activity of piperazine against Chikungunya virus targeting hydrophobic pocket of alphavirus capsid protein. Antiviral Research 146, 102–111. 10.1016/j.antiviral.2017.08.015

Angus, D.C., Berry, S., Lewis, R.J., Al-Beidh, F., Arabi, Y., van Bentum-Puijk, W., Bhimani, Z., Bonten, M., Broglio, K., Brunkhorst, F., Cheng, A.C., Chiche, J.D., de Jong, M., Detry, M., Goossens, H., Gordon, A., Green, C., Higgins, A.M., Hullegie, S.J., Kruger, P., Lamontagne, F., Litton, E., Marshall, J., McGlothlin, A., McGuinness, S., Mouncey, P., Murthy, S., Nichol, A., O’Neill, G.K., Parke, R., Parker, J., Rohde, G., Rowan, K., Turner, A., Young, P., Derde, L., McArthur, C., Webb, S.A., 2020. The remap-cap (Randomized embedded multifactorial adaptive platform for community-acquired pneumonia) Study rationale and design. Annals of the American Thoracic Society 17, 879–891. 10.1513/AnnalsATS.202003-192SD

Bhutkar, M., Singh, V., Dhaka, P., Tomar, S., 2022. Virus-host protein-protein interactions as molecular drug targets for arboviral infections. Frontiers in Virology 2, 959586. 10.3389/FVIRO.2022.959586

Blum, C.A., Roethlisberger, E.A., Cesana-Nigro, N., Winzeler, B., Rodondi, N., Blum, M.R., Briel, M., Mueller, B., Christ-Crain, M., Schuetz, P., 2023. Adjunct prednisone in community-acquired pneumonia: 180-day outcome of a multicentre, double-blind, randomized, placebo-controlled trial. BMC Pulmonary Medicine 23, 1–12. 10.1186/s12890-023-02794-w

Boniardi, I., Corona, A., Basquin, J., Basquin, C., Milia, J., Nagy, I., Tramontano, E., Zinzula, L., 2023. Suramin inhibits SARS-CoV-2 nucleocapsid phosphoprotein genome packaging function. Virus Research 336, 199221. 10.1016/j.virusres.2023.199221

Brady, D.K., Gurijala, A.R., Huang, L., Hussain, A.A., Lingan, A.L., Pembridge, O.G., Ratangee, B.A., Sealy, T.T., Vallone, K.T., Clements, T.P., 2024. A guide to COVID□19 antiviral therapeutics: a summary and perspective of the antiviral weapons against SARS□CoV□2 infection. The FEBS Journal 291, 1632–1662. 10.1111/febs.16662

Byun, W.G., Lim, D., Park, S.B., 2020. Discovery of Small□Molecule Modulators of Protein–RNA Interactions by Fluorescence Intensity□Based Binding Assay. ChemBioChem 21, 818–824. 10.1002/cbic.201900467

Caines, M.E.C., Bichel, K., Price, A.J., McEwan, W.A., Towers, G.J., Willett, B.J., Freund, S.M. V, James, L.C., 2012. Diverse HIV viruses are targeted by a conformationally dynamic antiviral. Nature Structural & Molecular Biology 19, 411–416. 10.1038/nsmb.2253

Cao, B., Wang, Y., Wen, D., Liu, W., Wang, Jingli, Fan, G., Ruan, L., Song, B., Cai, Y., Wei, M., Li, X., Xia, J., Chen, N., Xiang, J., Yu, T., Bai, T., Xie, X., Zhang, L., Li, C., Yuan, Y., Chen, H., Li, Huadong, Huang, H., Tu, S., Gong, F., Liu, Y., Wei, Y., Dong, C., Zhou, F., Gu, X., Xu, J., Liu, Z., Zhang, Y., Li, Hui, Shang, L., Wang, K., Li, K., Zhou, X., Dong, X., Qu, Z., Lu, S., Hu, X., Ruan, S., Luo, S., Wu, J., Peng, L., Cheng, F., Pan, L., Zou, J., Jia, C., Wang, Juan, Liu, X., Wang, S., Wu, X., Ge, Q., He, J., Zhan, H., Qiu, F., Guo, L., Huang, C., Jaki, T., Hayden, F.G., Horby, P.W., Zhang, D., Wang, C., 2020. A Trial of Lopinavir–Ritonavir in Adults Hospitalized with Severe Covid-19. New England Journal of Medicine 382, 1787–1799. 10.1056/NEJMoa2001282

Chakraborty, C., Sharma, A.R., Bhattacharya, M., Agoramoorthy, G., Lee, S.-S., 2021. The Drug Repurposing for COVID-19 Clinical Trials Provide Very Effective Therapeutic Combinations: Lessons Learned From Major Clinical Studies. Frontiers in Pharmacology 12, 704205. 10.3389/fphar.2021.704205

Chen, C.Y., Chang, C. ke, Chang, Y.W., Sue, S.C., Bai, H.I., Riang, L., Hsiao, C.D., Huang, T. huang, 2007. Structure of the SARS Coronavirus Nucleocapsid Protein RNA-binding Dimerization Domain Suggests a Mechanism for Helical Packaging of Viral RNA. Journal of Molecular Biology 368, 1075–1086. 10.1016/j.jmb.2007.02.069

Choudhary, S., Nehul, S., Kumar, K.A., Sharma, S., Rani, R., Saha, A., Sharma, G.K., Tomar, S., Kumar, P., 2022. Crystal structure and activity profiling of deubiquitinating inhibitors-bound to SARS-CoV-2 papain like protease revealed new allosteric sites for antiviral therapies. bioRxiv 2022.11.11.516107. 10.1101/2022.11.11.516107

Cubuk, J., Alston, J.J., Incicco, J.J., Singh, S., Stuchell-Brereton, M.D., Ward, M.D., Zimmerman, M.I., Vithani, N., Griffith, D., Wagoner, J.A., Bowman, G.R., Hall, K.B., Soranno, A., Holehouse, A.S., 2021. The SARS-CoV-2 nucleocapsid protein is dynamic, disordered, and phase separates with RNA. Nature Communications 12, 1936. 10.1038/s41467-021-21953-3

Dang, M., Song, J., 2022. CTD of SARS-CoV-2 N protein is a cryptic domain for binding ATP and nucleic acid that interplay in modulating phase separation. Protein Science 31, 345–356. 10.1002/pro.4221

Dhaka, P., Singh, A., Choudhary, S., Peddinti, R.K., Kumar, P., Sharma, G.K., Tomar, S., 2023. Mechanistic and thermodynamic characterization of antiviral inhibitors targeting nucleocapsid N-terminal domain of SARS-CoV-2. Archives of Biochemistry and Biophysics 750, 109820. 10.1016/j.abb.2023.109820

Edwards, P.M., 2002. Origin 7.0:□ Scientific Graphing and Data Analysis Software. Journal of Chemical Information and Computer Sciences 42, 1270. 10.1021/CI0255432

Emsley, P., Lohkamp, B., Scott, W.G., Cowtan, K., 2010. Features and development of Coot. Acta Crystallographica Section D Biological Crystallography 66, 486–501. 10.1107/S0907444910007493

Eslami, G., Mousaviasl, S., Radmanesh, E., Jelvay, S., Bitaraf, S., Simmons, B., Wentzel, H., Hill, A., Sadeghi, A., Freeman, J., Salmanzadeh, S., Esmaeilian, H., Mobarak, M., Tabibi, R., Jafari Kashi, A.H., Lotfi, Z., Talebzadeh, S.M., Wickramatillake, A., Momtazan, M., Hajizadeh Farsani, M., Marjani, S., Mobarak, S., 2020. The impact of sofosbuvir/daclatasvir or ribavirin in patients with severe COVID-19. Journal of Antimicrobial Chemotherapy 75, 3366–3372. 10.1093/jac/dkaa331

Fatma, B., Kumar, R., Singh, V.A., Nehul, S., Sharma, R., Kesari, P., Kuhn, R.J., Tomar, S., 2020. Alphavirus capsid protease inhibitors as potential antiviral agents for Chikungunya infection. Antiviral Research 179, 104808. 10.1016/j.antiviral.2020.104808

Gordon, D.E., Jang, G.M., Bouhaddou, M., Xu, J., Obernier, K., White, K.M., O’Meara, M.J., Rezelj, V. V., Guo, J.Z., Swaney, D.L., Tummino, T.A., Hüttenhain, R., Kaake, R.M., Richards, A.L., Tutuncuoglu, B., Foussard, H., Batra, J., Haas, K., Modak, M., Kim, M., Haas, P., Polacco, B.J., Braberg, H., Fabius, J.M., Eckhardt, M., Soucheray, M., Bennett, M.J., Cakir, M., McGregor, M.J., Li, Q., Meyer, B., Roesch, F., Vallet, T., Mac Kain, A., Miorin, L., Moreno, E., Naing, Z.Z.C., Zhou, Y., Peng, S., Shi, Y., Zhang, Z., Shen, W., Kirby, I.T., Melnyk, J.E., Chorba, J.S., Lou, K., Dai, S.A., Barrio-Hernandez, I., Memon, D., Hernandez-Armenta, C., Lyu, J., Mathy, C.J.P., Perica, T., Pilla, K.B., Ganesan, S.J., Saltzberg, D.J., Rakesh, R., Liu, X., Rosenthal, S.B., Calviello, L., Venkataramanan, S., Liboy-Lugo, J., Lin, Y., Huang, X.P., Liu, Y.F., Wankowicz, S.A., Bohn, M., Safari, M., Ugur, F.S., Koh, C., Savar, N.S., Tran, Q.D., Shengjuler, D., Fletcher, S.J., O’Neal, M.C., Cai, Y., Chang, J.C.J., Broadhurst, D.J., Klippsten, S., Sharp, P.P., Wenzell, N.A., Kuzuoglu-Ozturk, D., Wang, H.Y., Trenker, R., Young, J.M., Cavero, D.A., Hiatt, J., Roth, T.L., Rathore, U., Subramanian, A., Noack, J., Hubert, M., Stroud, R.M., Frankel, A.D., Rosenberg, O.S., Verba, K.A., Agard, D.A., Ott, M., Emerman, M., Jura, N., von Zastrow, M., Verdin, E., Ashworth, A., Schwartz, O., D’Enfert, C., Mukherjee, S., Jacobson, M., Malik, H.S., Fujimori, D.G., Ideker, T., Craik, C.S., Floor, S.N., Fraser, J.S., Gross, J.D., Sali, A., Roth, B.L., Ruggero, D., Taunton, J., Kortemme, T., Beltrao, P., Vignuzzi, M., García-Sastre, A., Shokat, K.M., Shoichet, B.K., Krogan, N.J., 2020. A SARS-CoV-2 protein interaction map reveals targets for drug repurposing. Nature 583, 459–468. 10.1038/s41586-020-2286-9

Günther, Sebastian, Reinke, P.Y.A., Fernández-Garciá, Y., Lieske, J., Lane, T.J., Ginn, H.M., Koua, F.H.M., Ehrt, C., Ewert, W., Oberthuer, D., Yefanov, O., Meier, S., Lorenzen, K., Krichel, B., Kopicki, J.D., Gelisio, L., Brehm, W., Dunkel, I., Seychell, B., Gieseler, H., Norton-Baker, B., Escudero-Pérez, B., Domaracky, M., Saouane, S., Tolstikova, A., White, T.A., Hänle, A., Groessler, M., Fleckenstein, H., Trost, F., Galchenkova, M., Gevorkov, Y., Li, C., Awel, S., Peck, A., Barthelmess, M., Schlünzen, F., Xavier, P.L., Werner, N., Andaleeb, H., Ullah, N., Falke, S., Srinivasan, V., Francą, B.A., Schwinzer, M., Brognaro, H., Rogers, C., Melo, D., Zaitseva-Doyle, J.J., Knoska, J., Penã-Murillo, G.E., Mashhour, A.R., Hennicke, V., Fischer, P., Hakanpää, J., Meyer, J., Gribbon, P., Ellinger, B., Kuzikov, M., Wolf, M., Beccari, A.R., Bourenkov, G., Stetten, D. Von, Pompidor, G., Bento, I., Panneerselvam, S., Karpics, I., Schneider, T.R., Garcia-Alai, M.M., Niebling, S., Günther, C., Schmidt, C., Schubert, R., Han, H., Boger, J., Monteiro, D.C.F., Zhang, L., Sun, X., Pletzer-Zelgert, J., Wollenhaupt, J., Feiler, C.G., Weiss, M.S., Schulz, E.C., Mehrabi, P., Karnǐcar, K., Usenik, A., Loboda, J., Tidow, H., Chari, A., Hilgenfeld, R., Uetrech, C., Cox, R., Zaliani, A., Beck, T., Rarey, M., Günther, Stephan, Turk, D., Hinrichs, W., Chapman, H.N., Pearson, A.R., Betzel, C., Meents, A., 2021. X-ray screening identifies active site and allosteric inhibitors of SARS-CoV-2 main protease. Science 372, 642–646. 10.1126/science.abf7945

Heskin, J., Pallett, S.J.C., Mughal, N., Davies, G.W., Moore, L.S.P., Rayment, M., Jones, R., 2022. Caution required with use of ritonavir-boosted PF-07321332 in COVID-19 management. The Lancet 399, 21–22. 10.1016/S0140-6736(21)02657-X

Hong, J.Y., Lin, S.C., Kehn-Hall, K., Zhang, K.M., Luo, S.Y., Wu, H.Y., Chang, S.Y., Hou, M.H., 2024. Targeting protein-protein interaction interfaces with antiviral N protein inhibitor in SARS-CoV-2. Biophysical journal 123, 478–488. 10.1016/J.BPJ.2024.01.013

Hu, X., Zhou, Z., Li, F., Xiao, Y., Wang, Z., Xu, J., Dong, F., Zheng, H., Yu, R., 2021. The study of antiviral drugs targeting SARS-CoV-2 nucleocapsid and spike proteins through large-scale compound repurposing. Heliyon 7, e06387. 10.1016/j.heliyon.2021.e06387

Jia, Z., Liu, C., Chen, Y., Jiang, H., Wang, Z., Yao, J., Yang, J., Zhu, J., Zhang, B., Yuchi, Z., 2022. Crystal structures of the SARS-CoV-2 nucleocapsid protein C-terminal domain and development of nucleocapsid-targeting nanobodies. FEBS Journal 289, 3813–3825. 10.1111/febs.16239

Kumar, R., Nehul, S., Singh, A., Tomar, S., 2021. Identification and evaluation of antiviral potential of thymoquinone, a natural compound targeting Chikungunya virus capsid protein. Virology 561, 36–46. 10.1016/j.virol.2021.05.013

Kumari, S., Mistry, H., Bihani, S.C., Mukherjee, S.P., Gupta, G.D., 2024. Unveiling potential inhibitors targeting the nucleocapsid protein of SARS-CoV-2: Structural insights into their binding sites. International Journal of Biological Macromolecules 273, 133167. 10.1016/j.ijbiomac.2024.133167

Lea, W.A., Simeonov, A., 2011. Fluorescence polarization assays in small molecule screening. Expert Opinion on Drug Discovery 6, 17–32. 10.1517/17460441.2011.537322

Lin, S.M., Lin, S.C., Hsu, J.N., Chang, C.K., Chien, C.M., Wang, Y.S., Wu, H.Y., Jeng, U.S., Kehn-Hall, K., Hou, M.H., 2020. Structure-Based Stabilization of Non-native Protein-Protein Interactions of Coronavirus Nucleocapsid Proteins in Antiviral Drug Design. Journal of Medicinal Chemistry 63, 3131–3141. 10.1021/acs.jmedchem.9b01913

Luan, X., Li, X., Li, Y., Su, G., Yin, W., Jiang, Y., Xu, N., Wang, F., Cheng, W., Jin, Y., Zhang, L., Xu, H.E., Xue, Y., Zhang, S., 2022. Antiviral drug design based on structural insights into the N-terminal domain and C-terminal domain of the SARS-CoV-2 nucleocapsid protein. Science Bulletin 67, 2327–2335. 10.1016/j.scib.2022.10.021

Malin, J.J., Suárez, I., Priesner, V., Fätkenheuer, G., Rybniker, J., 2020. Remdesivir against COVID-19 and Other Viral Diseases. Clinical Microbiology Reviews 34, 1–21. 10.1128/CMR.00162-20

McBride, R., Van Zyl, M., Fielding, B., 2014. The Coronavirus Nucleocapsid Is a Multifunctional Protein. Viruses 6, 2991–3018. 10.3390/v6082991

Mercaldi, G.F., Bezerra, E.H.S., Batista, F.A.H., Tonoli, C.C.C., Soprano, A.S., Shimizu, J.F., Nagai, A., da Silva, J.C., Filho, H.V.R., do Nascimento Faria, J., da Cunha, M.G., Zeri, A.C.M., Nascimento, A.F.Z., Proenca-Modena, J.L., Bajgelman, M.C., Rocco, S.A., Lopes-de-Oliveira, P.S., Cordeiro, A.T., Bruder, M., Marques, R.E., Sforça, M.L., Franchini, K.G., Benedetti, C.E., Figueira, A.C.M., Trivella, D.B.B., 2022. Discovery and structural characterization of chicoric acid as a SARS-CoV-2 nucleocapsid protein ligand and RNA binding disruptor. Scientific Reports 12, 18500. 10.1038/s41598-022-22576-4

Morse, M., Sefcikova, J., Rouzina, I., Beuning, P.J., Williams, M.C., 2023. Structural domains of SARS-CoV-2 nucleocapsid protein coordinate to compact long nucleic acid substrates. Nucleic Acids Research 51, 290–303. 10.1093/nar/gkac1179

Murshudov, G.N., Skubák, P., Lebedev, A.A., Pannu, N.S., Steiner, R.A., Nicholls, R.A., Winn, M.D., Long, F., Vagin, A.A., 2011. REFMAC5 for the refinement of macromolecular crystal structures. Acta Crystallographica Section D: Biological Crystallography 67, 355–367. 10.1107/S0907444911001314

Narayanan, A., Narwal, M., Majowicz, S.A., Varricchio, C., Toner, S.A., Ballatore, C., Brancale, A., Murakami, K.S., Jose, J., 2022. Identification of SARS-CoV-2 inhibitors targeting Mpro and PLpro using in-cell-protease assay. Communications Biology 5, 1–17. 10.1038/s42003-022-03090-9

Nguyen, T.H. Van, Lichière, J., Canard, B., Papageorgiou, N., Attoumani, S., Ferron, F., Coutard, B., 2019. Structure and oligomerization state of the C-terminal region of the Middle East respiratory syndrome coronavirus nucleoprotein. Acta Crystallographica Section D: Structural Biology 75, 8–15. 10.1107/S2059798318014948

Nicola, M., Alsafi, Z., Sohrabi, C., Kerwan, A., Al-Jabir, A., Iosifidis, C., Agha, M., Agha, R., 2020. The socio-economic implications of the coronavirus pandemic (COVID-19): A review. International Journal of Surgery 78, 185–193. 10.1016/j.ijsu.2020.04.018

Owen, D.R., Allerton, C.M.N., Anderson, A.S., Aschenbrenner, L., Avery, M., Berritt, S., Boras, B., Cardin, R.D., Carlo, A., Coffman, K.J., Dantonio, A., Di, L., Eng, H., Ferre, R., Gajiwala, K.S., Gibson, S.A., Greasley, S.E., Hurst, B.L., Kadar, E.P., Kalgutkar, A.S., Lee, J.C., Lee, J., Liu, W., Mason, S.W., Noell, S., Novak, J.J., Obach, R.S., Ogilvie, K., Patel, N.C., Pettersson, M., Rai, D.K., Reese, M.R., Sammons, M.F., Sathish, J.G., Singh, R.S.P., Steppan, C.M., Stewart, A.E., Tuttle, J.B., Updyke, L., Verhoest, P.R., Wei, L., Yang, Q., Zhu, Y., 2021. An oral SARS-CoV-2 M pro inhibitor clinical candidate for the treatment of COVID-19. Science 374, 1586–1593. 10.1126/science.abl4784

Panjikar, S., Parthasarathy, V., Lamzin, V.S., Weiss, M.S., Tucker, P.A., 2009. On the combination of molecular replacement and single-wavelength anomalous diffraction phasing for automated structure determination. Acta Crystallographica Section D Biological Crystallography 65, 1089–1097. 10.1107/S0907444909029643

Papageorgiou, N., Lichière, J., Baklouti, A., Ferron, F., Sévajol, M., Canard, B., Coutard, B., 2016. Structural characterization of the N-terminal part of the MERS-CoV nucleocapsid by X-ray diffraction and small-angle X-ray scattering. Acta Crystallographica Section D Structural Biology 72, 192–202. 10.1107/S2059798315024328

Peng, Y., Du, N., Lei, Y., Dorje, S., Qi, J., Luo, T., Gao, G.F., Song, H., 2020. Structures of the SARS □CoV□2 nucleocapsid and their perspectives for drug design. The EMBO Journal 39, e105938. 10.15252/embj.2020105938

Rafael Ciges-Tomas, J., Franco, M.L., Vilar, M., 2022. Identification of a guanine-specific pocket in the protein N of SARS-CoV-2. Communications biology 5, 711. 10.1038/s42003-022-03647-8

Rani, R., Long, S., Pareek, A., Dhaka, P., Singh, A., Kumar, P., McInerney, G., Tomar, S., 2022. Multi-target direct-acting SARS-CoV-2 antivirals against the nucleotide-binding pockets of virus-specific proteins. Virology 577, 1–15. 10.1016/j.virol.2022.08.008

Robert, X., Gouet, P., 2014. Deciphering key features in protein structures with the new ENDscript server. Nucleic Acids Research 42, W320–W324. 10.1093/NAR/GKU316

Rochel, N., Wurtz, J.M., Mitschler, A., Klaholz, B., Moras, D., 2000. The crystal structure of the nuclear receptor for vitamin D bound to its natural ligand. Molecular Cell 5, 173–179. 10.1016/S1097-2765(00)80413-X

Sabini, E., Ort, S., Monnerjahn, C., Konrad, M., Lavie, A., 2003. Structure of human dCK suggests strategies to improve anticancer and antiviral therapy. Nature Structural Biology 10, 513–519. 10.1038/nsb942

Sharma, R., Kesari, P., Kumar, P., Tomar, S., 2018. Structure-function insights into chikungunya virus capsid protein: Small molecules targeting capsid hydrophobic pocket. Virology 515, 223–234. 10.1016/j.virol.2017.12.020

Siddiqui, M.R., Alothman, Z.A., Wabaidur, S.M., 2014. Ultraperformance liquid chromatography-mass spectrometric method for determination of ampicillin and characterization of its forced degradation products. Journal of Chromatographic Science 52, 1273–1280. 10.1093/chromsci/bmt211

Singh, A., Ruchi, R., Pareek, A., Kumar, P., Tomar, S., 2023. In Silico Guided Drug Repurposing to Combat SARS-CoV-2 by Targeting Mpro, the Key Virus Specific Protease. ChemRxiv. 10.26434/chemrxiv-2023-qf6rb-v2

Sola, I., Almazán, F., Zúñiga, S., Enjuanes, L., 2015. Continuous and Discontinuous RNA Synthesis in Coronaviruses. Annual Review of Virology 2, 265–288. 10.1146/ANNUREV-VIROLOGY-100114-055218

Sola, I., Mateos-Gomez, P.A., Almazan, F., Zuñiga, S., Enjuanes, L., 2011. RNA-RNA and RNA-protein interactions in coronavirus replication and transcription. RNA Biology. 10.4161/hv.8.2.14991

Sturza, F., Gu□ă, □tefan D., Popescu, G.A., 2023. Antibiotics Used for COVID-19 In-Patients from an Infectious Disease Ward. Antibiotics 12, 150. 10.3390/antibiotics12010150

Szelazek, B., Kabala, W., Kus, K., Zdzalik, M., Twarda-Clapa, A., Golik, P., Burmistrz, M., Florek, D., Wladyka, B., Pyrc, K., Dubin, G., 2017. Structural Characterization of Human Coronavirus NL63 N Protein. Journal of Virology 91. 10.1128/JVI.02503-16

The CCP4 suite: Programs for protein crystallography, 1994. . Acta Crystallographica Section D: Biological Crystallography 50, 760–763. 10.1107/S0907444994003112

Wang, Y.T., Long, X.Y., Ding, X., Fan, S.R., Cai, J.Y., Yang, B.J., Zhang, X.F., Luo, R. hua, Yang, L., Ruan, T., Ren, J., Jing, C.X., Zheng, Y.T., Hao, X.J., Chen, D.Z., 2022. Novel nucleocapsid protein-targeting phenanthridine inhibitors of SARS-CoV-2. European Journal of Medicinal Chemistry 227, 113966. 10.1016/j.ejmech.2021.113966

Wei, X., Zhou, Y., Shen, X., Fan, L., Liu, D., Gao, X., Zhou, J., Wu, Y., Li, Y., Feng, W., Zhang, Z., 2024. Ciclopirox inhibits SARS-CoV-2 replication by promoting the degradation of the nucleocapsid protein. Acta Pharmaceutica Sinica B 14, 2505–2519. 10.1016/j.apsb.2024.03.009

Wu, C.H., Chen, P.J., Yeh, S. wei, 2014. Nucleocapsid Phosphorylation and RNA Helicase DDX1 Recruitment Enables Coronavirus Transition from Discontinuous to Continuous Transcription. Cell Host Microbe. 10.1016/j.chom.2014.09.009

Yan, W., Zheng, Y., Zeng, X., He, B., Cheng, W., 2022. Structural biology of SARS-CoV-2: open the door for novel therapies. Signal Transduction and Targeted Therapy. 10.1038/s41392-022-00884-5

Yang, M., He, S., Chen, X., Huang, Z., Zhou, Ziliang, Zhou, Zhechong, Chen, Q., Chen, S., Kang, S., 2021. Structural Insight Into the SARS-CoV-2 Nucleocapsid Protein C-Terminal Domain Reveals a Novel Recognition Mechanism for Viral Transcriptional Regulatory Sequences. Frontiers in Chemistry 8, 624765. 10.3389/fchem.2020.624765

Ye, Q., Lu, S., Corbett, K.D., 2021. Structural Basis for SARS-CoV-2 Nucleocapsid Protein Recognition by Single-Domain Antibodies. Frontiers in Immunology 12, 719037. 10.3389/fimmu.2021.719037

Ye, Q., West, A.M.V., Silletti, S., Corbett, K.D., 2020. Architecture and self-assembly of the SARS-CoV-2 nucleocapsid protein. Protein Science 29, 1890–1901. 10.1002/pro.3909

Zhang, Y., Wu, F., Han, Y., Wu, Y., Huang, L., Huang, Y., Yan, D., Jiang, X., Ma, J., Xu, W., 2024. Unraveling the assembly mechanism of SADS-CoV virus nucleocapsid protein: insights from RNA binding, dimerization, and epitope diversity profiling. Journal of Virology 98. 10.1128/jvi.00926-24

Zhou, R., Zeng, R., von Brunn, A., Lei, J., 2020. Structural characterization of the C-terminal domain of SARS-CoV-2 nucleocapsid protein. Molecular Biomedicine 1, 1–11. 10.1186/s43556-020-00001-4

Zinzula, L., Basquin, J., Bohn, S., Beck, F., Klumpe, S., Pfeifer, G., Nagy, I., Bracher, A., Hartl, F.U., Baumeister, W., 2021. High-resolution structure and biophysical characterization of the nucleocapsid phosphoprotein dimerization domain from the Covid-19 severe acute respiratory syndrome coronavirus 2. Biochemical and Biophysical Research Communications 538, 54–62. 10.1016/j.bbrc.2020.09.131

